## Supplementary file for "Gene expression plasticity followed by genetic change during colonization a high-elevation environment"

She et al.

Figure S1. WGCNA identified co-expressed genes with gene expression change association with the three stages (*i.e.*, ancestral stage, plastic stage and colonized stage). (a)-(b) The soft thresholds of the scale independence used for WGCNA analyses were 23 and 4 for flight and cardiac muscles. (c)-(d) WGCNA identified six and nine highly correlated modules for the flight muscle (left) and cardiac muscle (right) transcriptomes using a threshold of $P$<0.1.

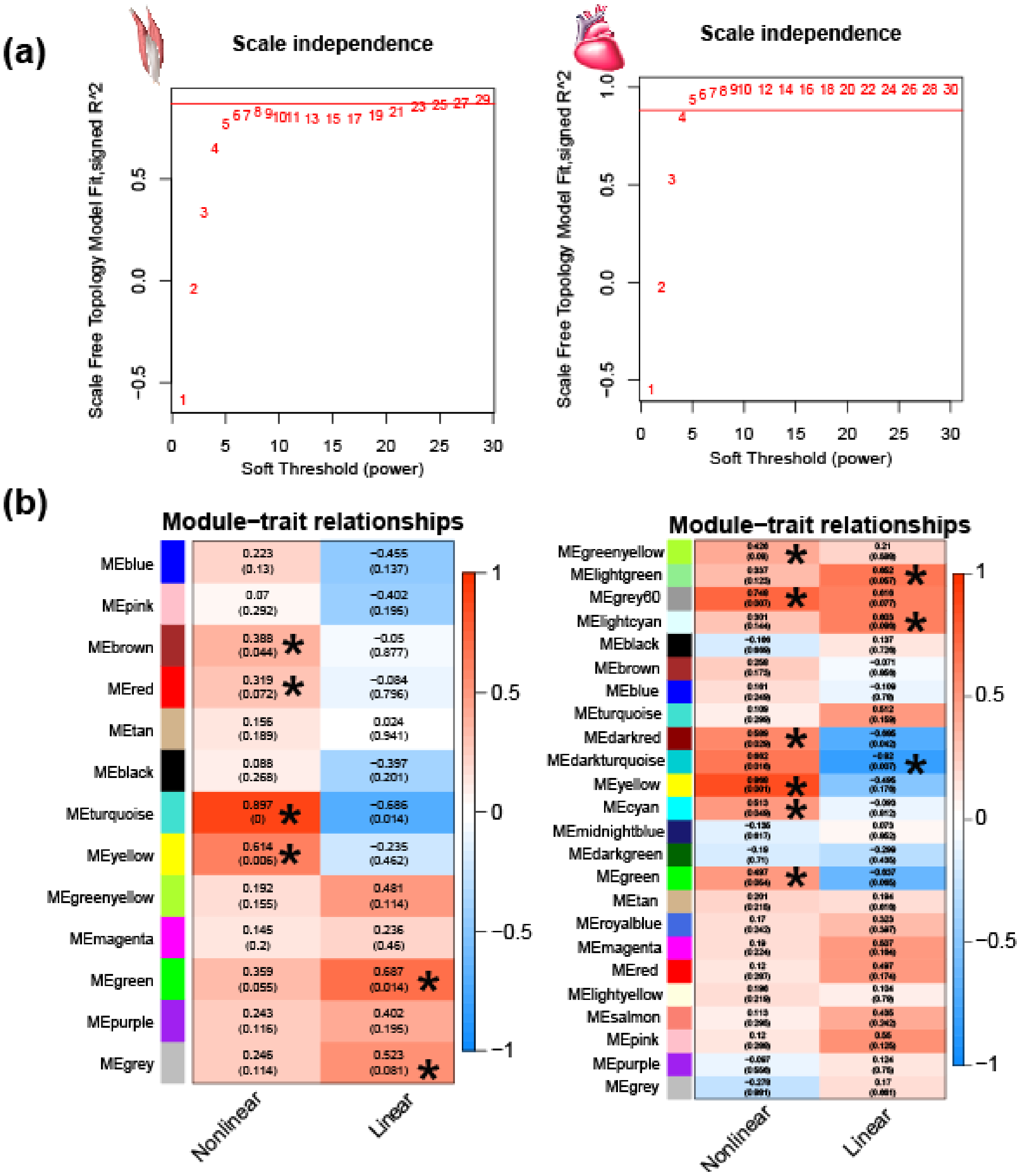

Supplementary Figure S2. Frequencies of genes with reinforcement and reversion plasticity in the flight and cardiac muscles. Three different bins are set along the spectrum of the reinforcement/reversion plasticity, *i.e.*, 20%, 40% and 60%. All comparisons show that there are more genes showing reversion plasticity than those showing reinforcement plasticity. Upper, flight muscle; lower, cardiac muscle. Two-tailed binomial test, ***, $P < 0.001$.

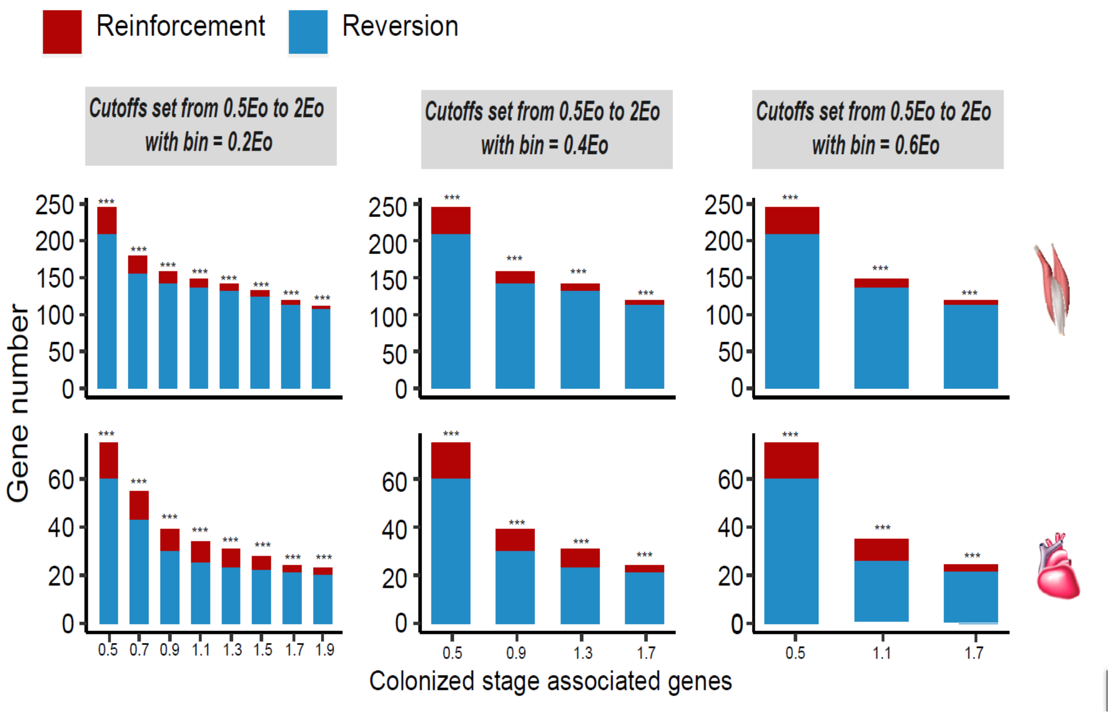

Supplementary Figure S3. Pearson correlation between gene expression and muscle phenotypic values of the ancestral and colonized individuals. (a) Red or blue dots showed the genes with their expression changes that are positively or negatively correlated with muscle phenotypes, respectively. A threshold of $P<0.05$ (vertical grey line) was used to determine statistical significance. (b) 2037 and 1866 muscle phenotype-associated genes were identified in the flight and cardiac muscle transcriptomes, respectively, which included 328 and 843 genes that are positively correlated with flight and cardiac muscle phenotypes, and 1709 and 1023 negatively correlated with flight and cardiac muscle phenotypes.

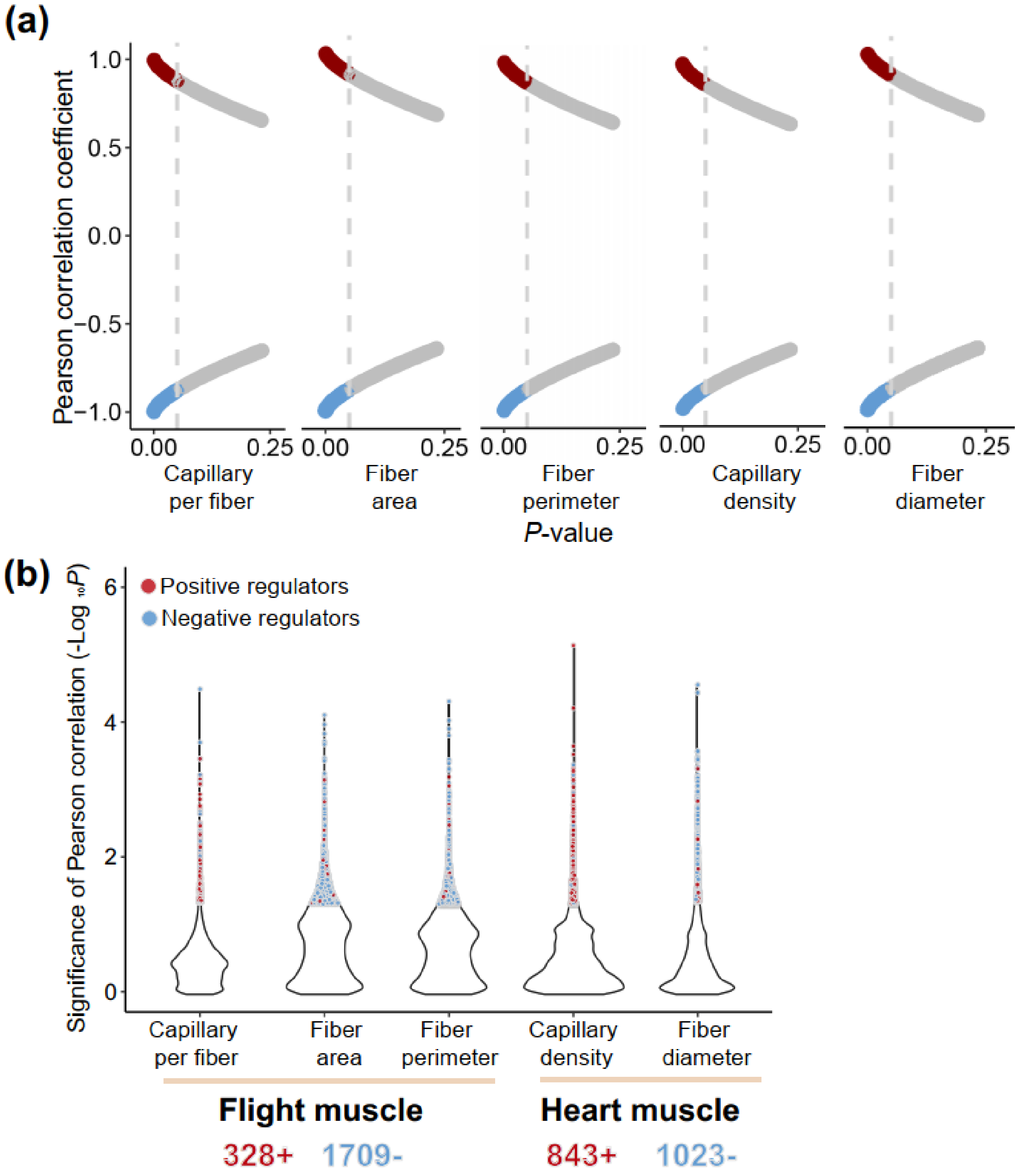

Supplementary Figure S4. Frequencies of genes with adaptive and maladaptive plasticity in the flight and cardiac muscles. Three different bins are set along the spectrum of the maladaptive/adaptive plasticity, *i.e.*, 20%, 40% and 60%. All comparisons show that there are more genes showing maladaptive plasticity than those showing adaptive plasticity for the flight muscle, while more than 50% of comparisons supported an excess of genes with maladaptive plasticity for the cardiac muscle. Two-tailed binomial test, NS, non significant; *, $P < 0.05$; **, $P < 0.01$; ***, $P < 0.001$.

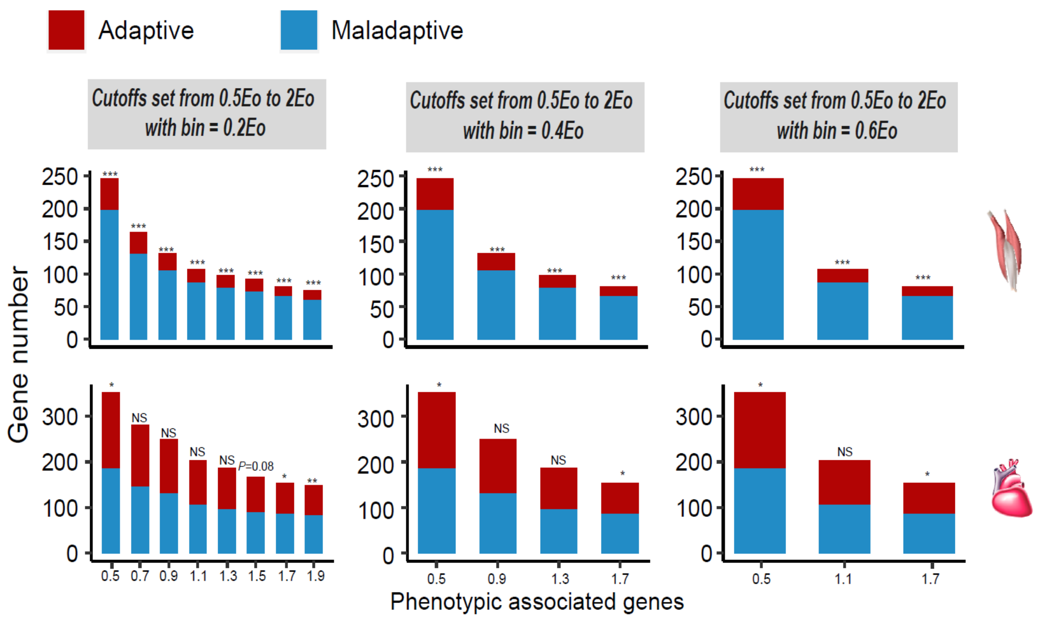

Supplementary Figure 5. The median expression levels of the conserved genes (*i.e.*, coefficient of variance ≤ 0.3 and average TPM ≥ 1 for each sample) did not differ among the lowland, hypoxia-exposed lowland and highland tree sparrows (Wilcoxon signed-rank test, $P$<0.05).

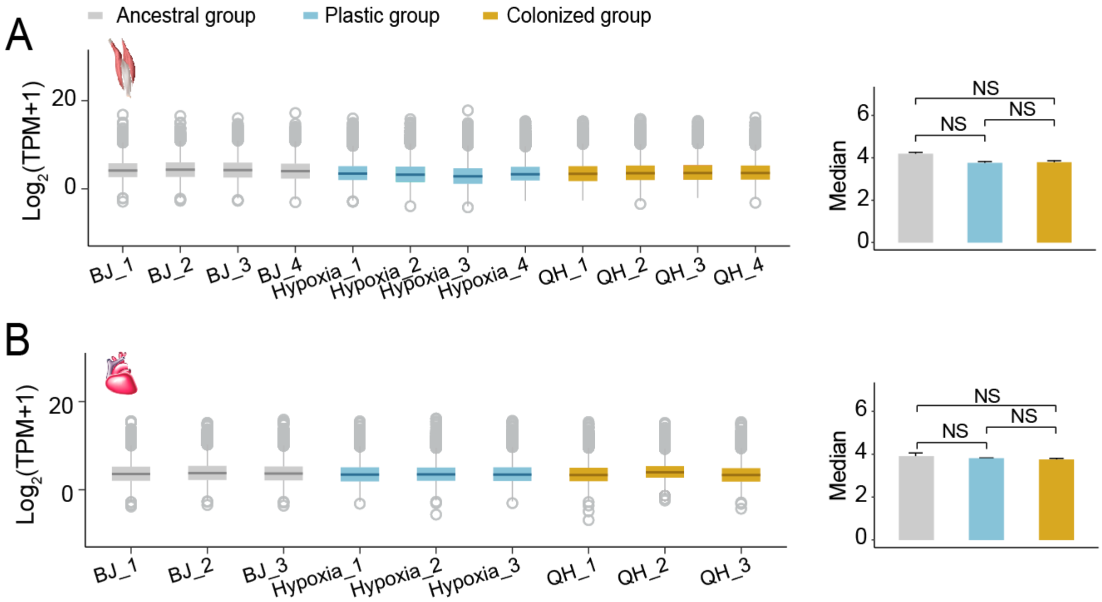

Supplementary Table S1. Sampling of transcriptomic data used in this study. All birds collected were adults and at pre-breeding season. Sex was unknown.

| Tissue | Groups | Individual ID | Sampling locality | Clean reads | Mapped reads | Percent mapped to genome (%) |
|---|---|---|---|---|---|---|
| Flight muscle | Ancestral Stage | BJ_1 | Beijing | 17,166,225 | 15,504,534 | 90.32% |
| | Ancestral Stage | BJ_2 | Beijing | 20,441,592 | 18,002,910 | 88.07% |
| | Ancestral Stage | BJ_3 | Beijing | 16,679,631 | 14,457,904 | 86.68% |
| | Ancestral Stage | BJ_4 | Beijing | 20,923,512 | 17,881,233 | 85.46% |
| | Colonized stage | QH_1 | Qinghai Lake | 16,690,586 | 14,589,241 | 87.41% |
| | Colonized stage | QH_2 | Qinghai Lake | 17,278,802 | 15,141,414 | 87.63% |
| | Colonized stage | QH_3 | Qinghai Lake | 16,849,769 | 14,802,522 | 87.85% |
| | Colonized stage | QH_4 | Qinghai Lake | 17,144,248 | 14,857,205 | 86.66% |
| | Plastic stage | Hypoxia_1 | Beijing | 16,937,478 | 14,754,237 | 87.11% |
| | Plastic stage | Hypoxia_2 | Beijing | 18,632,409 | 16,465,460 | 88.37% |
| | Plastic stage | Hypoxia_3 | Beijing | 20,693,777 | 18,202,246 | 87.96% |
| | Plastic stage | Hypoxia_4 | Beijing | 20,686,863 | 17,995,502 | 86.99% |
| Cardiac muscle | Ancestral stage | BJ_1 | Beijing | 21,447,101 | 19,109,367 | 89.10% |
| | Ancestral stage | BJ_2 | Beijing | 17,223,560 | 15,299,688 | 88.83% |
| | Ancestral stage | BJ_3 | Beijing | 20,244,087 | 17,636,649 | 87.12% |
| | Colonized stage | QH_1 | Qinghai Lake | 17,350,254 | 15,256,078 | 87.93% |
| | Colonized stage | QH_2 | Qinghai Lake | 17,116,185 | 15,087,917 | 88.15% |
| | Colonized stage | QH_3 | Qinghai Lake | 17,288,719 | 15,504,523 | 89.68% |
| | Plastic stage | Hypoxia_1 | Beijing | 17,120,796 | 14,960,152 | 87.38% |
| | Plastic stage | Hypoxia_2 | Beijing | 16,697,228 | 14,757,010 | 88.38% |
| | Plastic stage | Hypoxia_3 | Beijing | 16,859,063 | 15,166,413 | 89.96% |

**Supplementary Table 2.** Statistics of reads mapping and coverage of 11 highland and 12 lowland tree sparrows (Qu et al. 2020). All birds collected were adults and sex is unknown.

| Group | Sample | Location | Collection time | Longitude | Latitude | Elevation | Clean reads (G) | Coverage | Mapping rate (%) |
|---|---|---|---|---|---|---|---|---|---|
| Highland | IOZ14015 | Gangcha | 2014-4-10 | 99.74 | N37.03 | 3213m | 17.94 | 15.40 | 98.80 |
| | IOZ14018 | Gangcha | 2014-4-11 | E99.74 | N37.03 | 3213m | 15.52 | 13.15 | 98.55 |
| | IOZ14019 | Gangcha | 2014-4-11 | E99.74 | N37.03 | 3213m | 18.13 | 15.34 | 98.75 |
| | IOZ14020 | Gangcha | 2014-4-11 | E99.74 | N37.03 | 3213m | 18.24 | 15.66 | 98.77 |
| | IOZ14021 | Gangcha | 2014-4-11 | E99.74 | N37.03 | 3213m | 18.24 | 15.66 | 98.77 |
| | IOZ6498 | Qinghai lake | 2008-2-26 | E99.47 | N36.59 | 3200m | 16.45 | 14.00 | 98.80 |
| | IOZ6499 | Qinghai lake | 2008-2-26 | E99.47 | N36.59 | 3200m | 14.19 | 12.07 | 98.79 |
| | IOZ6500 | Qinghai lake | 2008-2-26 | E99.47 | N36.59 | 3200m | 14.72 | 12.50 | 98.86 |
| | IOZ6501 | Qinghai lake | 2008-2-26 | E99.47 | N36.59 | 3200m | 16.85 | 14.27 | 98.66 |
| | IOZ6502 | Qinghai lake | 2008-2-26 | E99.47 | N36.59 | 3200m | 18.89 | 16.03 | 98.75 |
| | IOZ6503 | Qinghai lakemahe | 2008-2-26 | E99.47 | N36.59 | 3200m | 23.13 | 19.51 | 98.86 |
| Lowland | IOZ4735 | Qinghuangdao | 2010-11-6 | E119.35 | N39.56 | 100m | 21.79 | 18.59 | 98.79 |
| | IOZ4736 | Qinghuangdao | 2010-11-6 | E119.35 | N39.56 | 100m | 20.83 | 17.81 | 98.93 |
| | IOZ4737 | Qinghuangdao | 2010-11-6 | E119.35 | N39.56 | 100m | 19.86 | 16.75 | 98.86 |
| | IOZ4738 | Qinghuangdao | 2010-11-6 | E119.35 | N39.56 | 100m | 22.32 | 19.01 | 94.66 |
| | IOZ4739 | Qinghuangdao | 2010-11-6 | E119.35 | N39.56 | 100m | 20.13 | 17.08 | 98.90 |
| | IOZ4740 | Qinghuangdao | 2010-11-6 | E119.35 | N39.56 | 100m | 28.55 | 24.24 | 98.76 |
| | IOZ14001 | Bejing | 2014-3-24 | E116.35 | N39.93 | 59m | 19.15 | 16.16 | 98.77 |
| | IOZ14002 | Bejing | 2014-3-24 | E116.35 | N39.93 | 59m | 20.32 | 17.27 | 98.80 |
| | IOZ14003 | Bejing | 2014-3-24 | E116.35 | N39.93 | 59m | 20.75 | 17.74 | 98.73 |
| | IOZ14004 | Bejing | 2014-3-24 | E116.35 | N39.93 | 59m | 28.88 | 24.31 | 98.76 |
| | IOZ14006 | Tianjin | Unknown | E117.10 | N39.10 | 70m | 19.71 | 16.65 | 98.67 |
| | IOZ18809 | Tianjin | Unknown | E117.10 | N39.10 | | 33.53 | 28.44 | 98.72 |
